# Hypertonic lactate infusion reduces vasopressor requirement and biomarkers of brain and cardiac injury after experimental cardiac arrest

**DOI:** 10.1101/2023.03.07.531627

**Authors:** Filippo Annoni, Fuhong Su, Lorenzo Peluso, Ilaria Lisi, Enrico Caruso, Francesca Pischiutta, Elisa Gouvea Bogossian, Bruno Garcia, Hassane Njimi, Jean-Louis Vincent, Nicolas Gaspard, Lorenzo Ferlini, Jacques Creteur, Elisa R Zanier, Fabio Silvio Taccone

## Abstract

**Introduction:** Prognosis after resuscitated cardiac arrest (CA) remains poor with high morbidity and mortality due to extensive cardiac and brain injuries and the lack of effective treatments. Hypertonic sodium lactate (HSL) could be beneficial after CA by buffering severe metabolic acidosis, increasing brain perfusion and cardiac performance, reducing cerebral swelling, and serving as alternative energetic cellular substrate. The aim of this study was therefore to test the effects of HSL infusion on brain and cardiac injury in an experimental model of CA.

**Methods:** After a 10-min electrically induced CA followed by 5 min of cardiopulmonary resuscitation maneuvers, adult swine (n=35) were randomly assigned to receive either balanced crystalloids (controls, n=11) or HSL infusion, either starting during cardiopulmonary resuscitation (CPR, Intra-arrest, n=12) or after return of spontaneous circulation (Post-ROSC, n=11) for the following 12 hours. In all animals, extensive multimodal neurological and cardiovascular monitoring was implemented. All animals were treated with target temperature management at 34°C.

**Results:** 34 out of 35 (97.1%) animals achieved ROSC and one animal in the Intra-arrest group deceased before completing the observation period. Arterial pH, lactate, sodium concentrations and plasma osmolarity were higher in treated animals then in controls (p<0.001), while potassium concentrations were lower (p=0.004). HSL infusion either Intra-arrest or Post-ROSC improved hemodynamic compared to controls, as shown by reduced vasopressors need to maintain mean arterial pressure target above 65 mmHg (p=0.005 for interaction; p=0.01 for groups). Moreover, plasmatic troponin-I levels and glial fibrillary acid protein (GFAP) concentrations were lower in treated groups at several time-points than in controls.

**Conclusions:** In this experimental CA model, HSL infusion was associated with reduced vasopressor requirements and decreased plasmatic levels of biomarkers of cardiac and cerebral injury.

## Introduction

Sudden cardiac arrest (CA) remains a prominent health problem worldwide, being a leading cause of morbidity and mortality and a massive healthcare burden [1]. Even when cardiopulmonary resuscitation (CPR) succeeds and return of spontaneous circulation (ROSC) is achieved, survivors remain at high-risk of death in the following days and only a minority of them returns to a quality of life similar to prior the event [2]. Among patients admitted to the hospital after resuscitated CA, extensive cardiac and brain injuries represent the leading cause of unfavorable outcomes, the latter accounting for most deaths [3]. Several drugs and interventions have been tried unsuccessfully [4]. Even target temperature management has not been proven to significantly improve neurological outcome of CA patients [5–7].

In traumatic brain injury patients, hypertonic sodium lactate (HSL) has been proven safe and capable to reduce intracranial pressure (ICP), prevent episodes of raised ICP, improve cerebral metabolism and determines a cerebral glucose sparing effect [8–13]. Moreover, HSL infusion could improve cardiac performance in some selected populations, such as patients with acute heart failure [15–16]. As such, HSL infusion could represent a suitable therapeutic option in CA patients; in particular, HSL could: 1) provide alternative energetic substrate for neurons in situation of metabolic stress; 2) reduce glutamate-related excitotoxicity; 3) mitigate severe post-resuscitation metabolic acidemia; 4) decrease intracranial pressure and cellular swelling; 5) improve post-resuscitation myocardial dysfunction [14].

HSL administration has only been studied in one rabbit model of CA, where it increased mean arterial pressure and improved cardiac function and cerebral mitochondrial function and reduced brain injury as expressed by plasmatic levels of the protein S100ß [17]. Large animal models are best suited for research in CA, allowing the usage of mechanical CPR, frequent blood sampling and invasive monitoring devices. Furthermore, swine have a gyrencephalic brain architecture and cardiovascular system that closely resemble to the human one, increasing the possibility for translatability of experimental findings.

The aim of this study was therefore to assess the effects of HSL infusion during and after CPR on the cardiovascular system, on brain function and perfusion and on related biomarkers of organ injury.

## Methods

### Experimental procedure

The Institutional Review Board for Animal Care of the Free University of Brussels (Belgium) approved all experimental procedures (number of Ethical Committee approval: 704 N), which were also in compliance with ARRIVE (Animal Research: Reporting in Vivo Experiments) guidelines. Care and handling of the animals were in accord with National Institutes of Health guidelines (Institute of Laboratory Animal Resources).

A detailed description of this experimental model has been previously published [19], and supplementary information regarding animal handling are provided in the Supplementary Material. Briefly, on the day of the experiment the animal (adult swine, both sexes) was initially sedated in the cage and transported to the operating theater where, after being equipped with an arterial catheter in the femoral artery and peripheral venous catheter, was intubated and sedated with a 1% mixture of inhaled sevoflurane. The animal was then equipped with a Foley catheter, a three-lumen central venous line in the external jugular vein, a pulmonary artery catheter and a single lumen central venous catheter inserted upstream to allow sampling of brain effluent blood. After pronation, multimodal neuromonitoring was placed surgically and included two cerebral microdialysis (CMD) catheters (one in each frontal lobe), a single probe measuring ICP, cerebral temperature and brain tissue oxygen tension (PbtO_2_) in one parietal lobe, a laser Doppler probe in the other parietal lobe and one stereoelectroencephalography (sEEG) wire in each parietal lobe. After stabilization, ventricular fibrillation was induced electrically via a pacing wire and left untreated for 10 min. After this period, chest compressions were started at the rate of 100/min for 5 min and ventilation was resumed with a FiO_2_ of 1, along with sedation and analgesia. After one minute, an intravenous injection of epinephrine was administered. At the end of the five minutes window, a biphasic electric countershock was delivered. CPR was eventually restarted for and additional minute in case of unsuccessful shock and another countershock was delivered every minute along with a second dose of epinephrine for a total CPR time of seven min. ROSC was considered achieved if the rhythm remained compatible with a mean arterial pressure (MAP) above 65 mmHg for at least 20 min. If the animal failed to achieve ROSC after 12 shocks, it was considered dead. If ROSC was achieved, the animal was proned again and observed for 12 additional hours. Death was confirmed by ventricular fibrillation waves on the EKG trace and the concomitant drop of arterial blood pressure. All animals received target temperature management aiming at a temperature of 34C°.

### Group allocation and treatment preparation

On the day of the experiment, animals were randomly assigned (simple randomization) to receive either a bolus of 20 mL of NaCl 0.9% at the beginning of CPR maneuvers and a continuous infusion of balanced crystalloids during the observation period (Control group) or an intravenous bolus of 10 mmoL of HSL (Monico SpA, Venezia, Italy) diluted in 20 mL of NaCl 0.9% at CPR initiation followed by a continuous infusion of 30 μmol/Kg/min during the observation period (Intra-arrest group) or 20 mL of NaCl 0.9% at the beginning of CPR followed by a continuous infusion of 30 μmol/Kg/min HSL during the observation period (Post-ROSC group, **Figure 1**). Due to the magnitude of the changes in pH and lactate, blinding was not feasible. In the treated groups, specific safety limits were pre-established to standardize the administration of HSL and reproduce conditions similar to clinical practice. Each hour, according to the arterial blood gas analysis, the HSL perfusion rate was reduced by 20% if any of the following criteria were met: arterial pH increase >0.10 compared to baseline pH or >7.65; Na^+^ ≥155 mEq/l or plasmatic Osm ≥ 320 mOsm.

**Figure 1:**
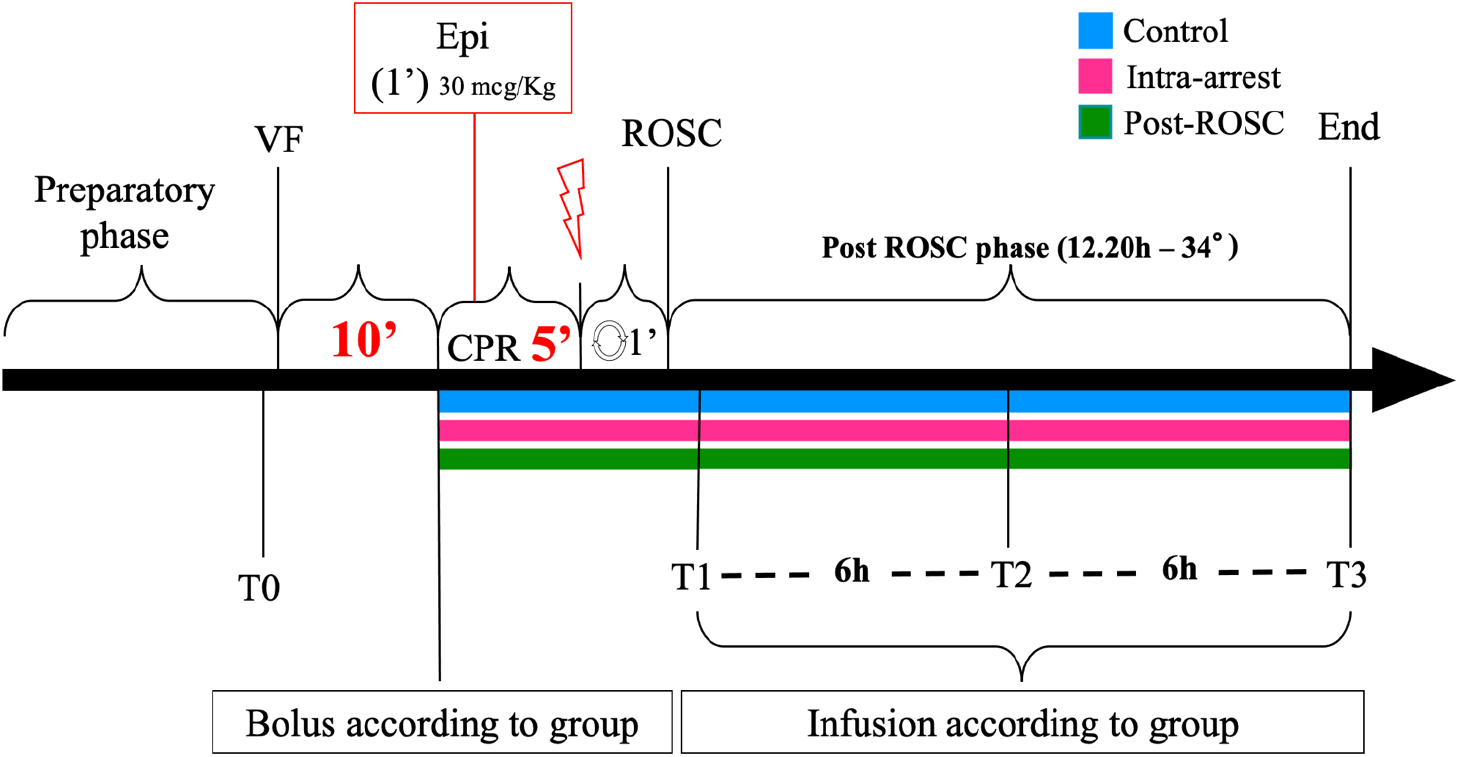
Timeline of the experiment. Control group is represented in blue, Intra-arrest group in pink and Post-ROSC group in green. VF: ventricular fibrillation; Epi: epinephrine; CPR: cardiopulmonary resuscitation; ROSC: return of spontaneous circulation; T0-3: blood sampling timepoints.

During the whole experiment, a minimal MAP of at least 65 mmHg was maintained with norepinephrine. All animals received a fixed continuous infusion of balanced crystalloids (5-10 ml/kg/h) and any additional fluid administration was titrated to keep the value of pulse pressure variation (PPV) < 14%.

Hyperglycemia after ROSC was left uncorrected if < 300 mg/dl within the first hour from ROSC and < 250 mg/dl during the study period and otherwise corrected with the infusion of 10U of regular insulin (Actrapid, Novo Nordisk, Bagsværd, Denmark).

### Blood and brain interstitial fluid sampling

Arterial blood samples were obtained prior to CA induction (T0), 20 min after ROSC (sustained ROSC, T1), and 6 hours (T2) and 12 hours later (T3), centrifugated and separated for plasma to be stored (10 mL) at −80°C. Arterial and jugular blood gas analyses were carried out before and after endotracheal intubation, at T0 and T1, and then hourly until the end of the experiment, and arterio-jugular difference for lactate and glucose were computed at each time point, whereas central venous mixed blood was collected at baseline, T1 and every three hours after to allow instrument calibration. CMD sampling was obtained at T0 and every hour after T1 from the same catheter. The spare CMD sample was stored at −80°C in case of necessity.

### Brain tissue collection

After the animal sacrifice, the frontal and parietal bones were carefully opened with the help of a Kerrison bone punch, the dura mater was dissected with a scalp and one section of the parietal cortex of approximately 0.5 cm^3^ was harvested in each side and immediately frozen in liquid nitrogen and stored at −80°C.

### Gene expression analysis

Total RNA was extracted from the cortex with a Ribospin II kit (GeneAll, #314-103). Samples were reverse transcribed with random hexamer primers using Multi-Scribe Reverse Transcriptase (Life Technologies). Real-time reverse transcription PCR was done with *RPL27* as housekeeping gene. Relative gene expression was determined by ΔΔCt method. Data are expressed as the log2-fold difference over the sham adult group. Genes and primer sequences are listed in **Table S2**. The explorative analysis included the following genes: Microtubule-Associated Protein 2 (MAP2); Glial Fibrillary Acid Protein (GFAP); Cluster of Differentiation molecule 11ß (CD11ß); Platelet and Endothelial Cell Adhesion Molecule 1 (PECAM1); Caspase 3 and 8 (CASP3 and CASP8) and Heme Oxygenase 1 (HO-1).

### Brain injury biomarkers

Plasma levels of GFAP, neurofilament light chain protein (NFL) and neuron specific enolase (NSE) were measured using commercially available single molecule array assay kits on an SR-X Analyzer: GFAP (#102336), NFL (#103400) and NSE (#102475) as described by the manufacturer (Quanterix, Billerica, MA). A single batch of reagents was used for each analyte.

### Continuous intracerebral EEG monitoring

All sEEG traces acquired via the dedicated device and transferred to the acquisition software were analyzed considering a 10-min window for baseline prior to the beginning of cardiac arrest procedure (T0). During the CA procedure (i.e., prone position, no-flow time, low-flow time and countershock) the sEEG electrodes were disconnected to prevent electrical damage of the experimental tools from the countershock and reconnected as soon as possible after ROSC. The recording lasted during the entire observation phase (i.e., 12 hours). The EEG signal with the best signal-to-noise ratio was selected for further analysis. All analyses were performed offline, using built-in and custom functions in Matlab. We filtered the EEG signal between 1 and 15 Hz (4th order butterworth bandpass filter, filtfilt function in Matlab), and then we computed the Hilbert transform (Hilbert function in Matlab) of the filtered signal and extracted the amplitude. We calculated the mean, kurtosis, skewness, and standard deviation of the amplitude using a sliding 1 minute window with 50% overlap. Further, the EEG background at T3 [20] was assessed for the presence of a suppressed background or burst suppression by two independent neurophysiologists, expert in EEG reading and blinded to the study assignment.

### Data handling and reduction

Multiple variables were recorded continuously with a sampling frequency between 1 and 100 Hz, including blood pressure (systolic, diastolic, mean), heart rate, ICP, PbtO_2_, brain temperature and cerebral blood flow (CBF). Data were extracted as means over period of 60 seconds and successively reduced to means over 10 minutes. For the sEEG signal, the amplitude of the Hilbert transform of the filtered signal between 1 and 15 Hz was computed. Outliers were detected and eliminated using the ROUT method with a Q = 1% (where Q is the maximum desired false discovery rate).

### Statistical Analysis

With an expected survival rate of >90% [20] and given the absence of prior studies on the subject, an *a priori* sample size of 10 animals per group was considered adequate for the study purposes. Continuous variables are expressed as means with standard deviation or medians with interquartile range, accordingly to the population’s distribution, and discrete variables are expressed in percentages with 95% confidence intervals. Categorical variables were compared with Fisher’s exact test or Chi-square test, as appropriate. For multiple group comparison, ANOVA test, Welch’s test, Kruskal-Wallis test or Dunnett’s test were used as appropriate. To compare the time-based variations between groups, a linear mixed-effect models fitted for restricted maximum likelihood estimation (REML) was used. Two different analyses were conducted for cerebral lactate and LPR (between T0 and T2 and between T2 and T3), as well as in the EEG variables (between T1 and T2 and between T2 and T3). For variation in gene expression compared to the Control group, one sample t test or Wilcoxon test were used, as appropriate. A value of p<0.05 was considered statistically significant. Data analyses were performed with GraphPad Prism (version 9.3.1 for Macintosh, GraphPad Software, La Jolla, CA, US) and Matlab (version R2019b, The MathWorks Inc., Natick, MA, US).

## Results

Of the 35 animals that underwent the CA procedure, 34 (97.1%) achieved ROSC and were included in the study, 11 in the Control group, 12 in the Intra-arrest group and 11 in the Post-ROSC group. One animal in the Intra-arrest group had a secondary VF episode 7 hours after ROSC, possibly related to the mobilization of the pulmonary catheter: despite a successful countershock, data acquired beyond this timepoint, as well as the brain tissue samples, were excluded from analysis.

There was no significant difference between the three groups in weight and sex, time to target core temperature of 34°C (3.9 ± 0.6 hours; p=0.89) and HSL infusion rate in the treated groups (**Figure 2**). In both Intra-arrest and Post-ROSC groups, we assisted to a similar decrease in HSL infusion rate from the fifth hour post ROSC, according to the pre-specified safety protocol. A more detailed description of baseline characteristics of the study groups is reported in **Table 1**.

**Figure 2:**
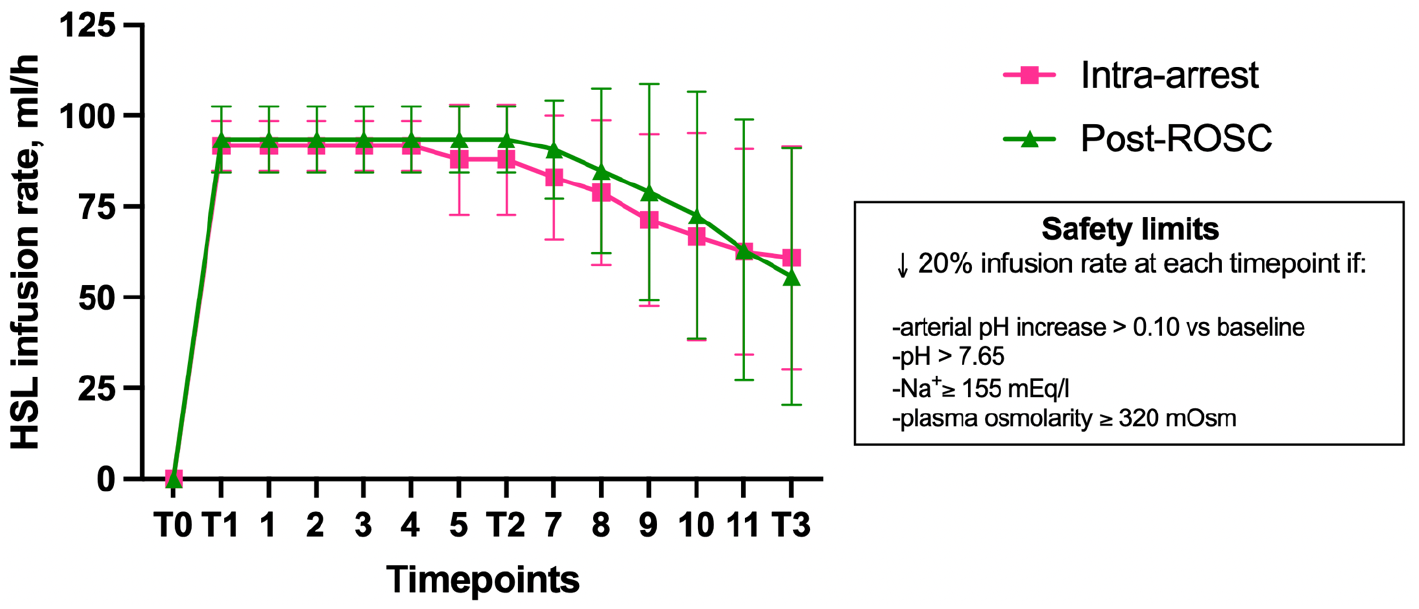
HSL infusion rate in treated groups. (Means ± SD)

**Table 1:**
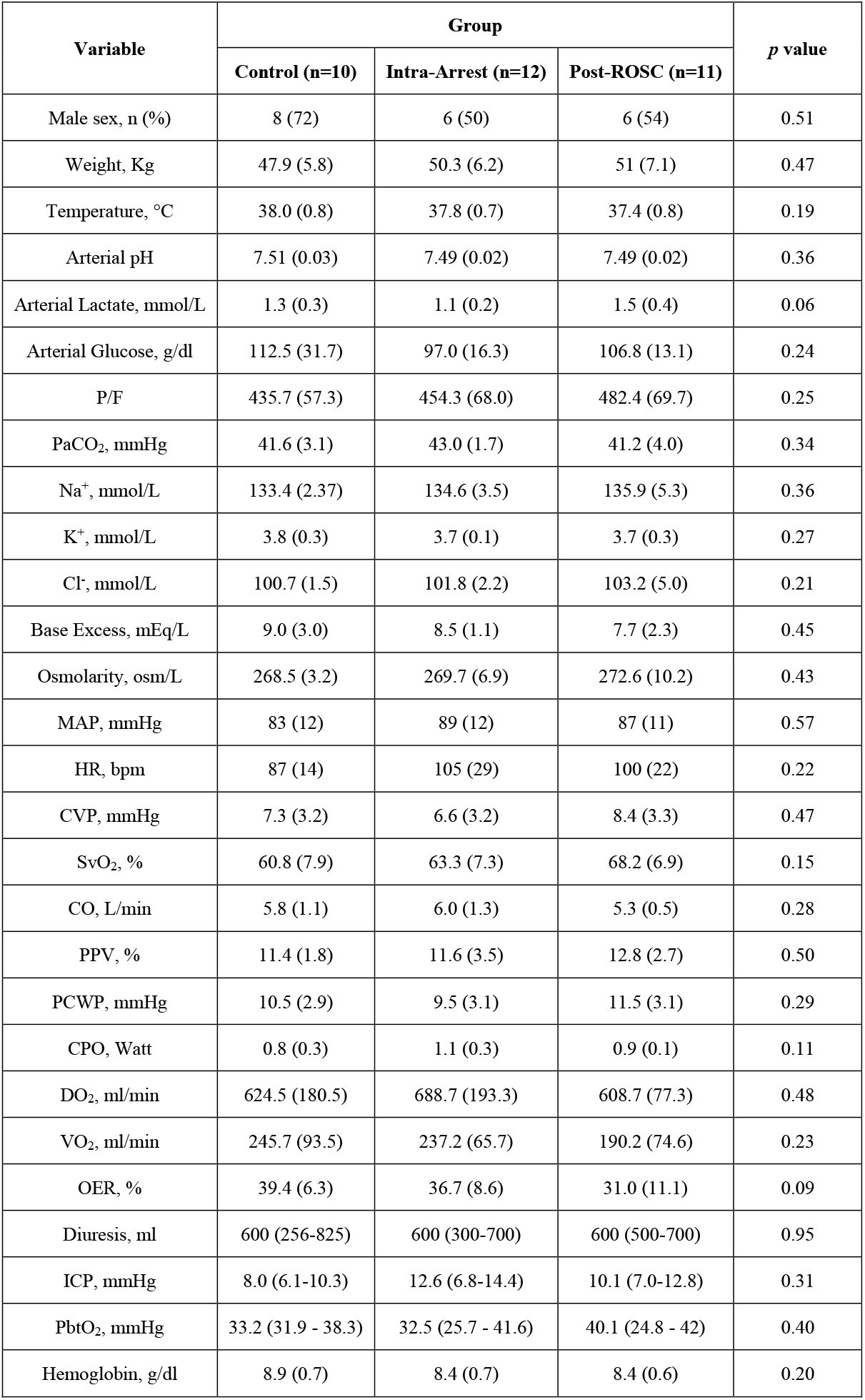
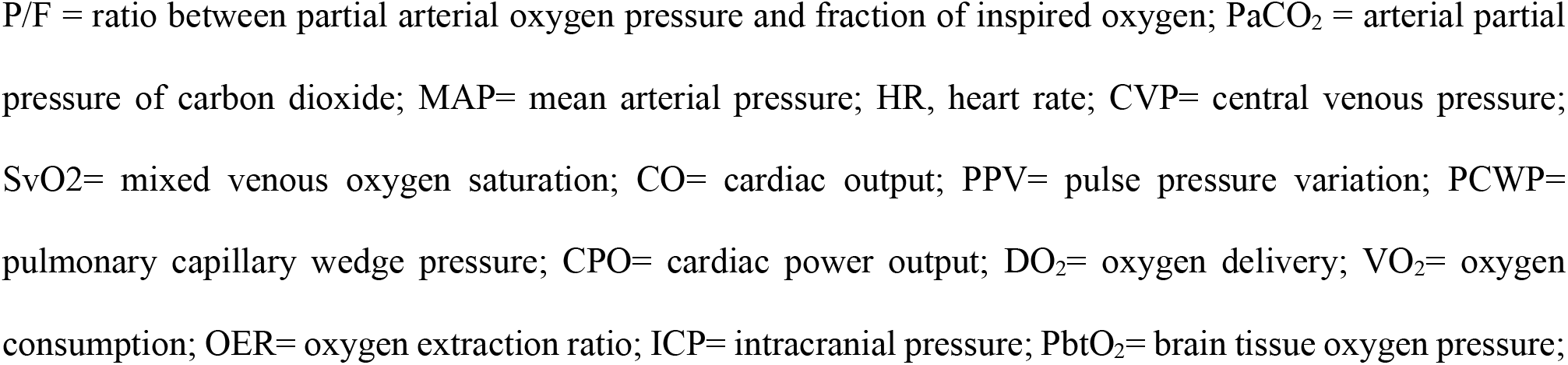
Baseline characteristics of the study groups. Data are reported as mean (SD).

| Variable | Group |  |  | *p* value |
| --- | --- | --- | --- | --- |
|  | Control (n=10) | Intra-Arrest (n=12) | Post-ROSC (n=11) |  |
| Male sex, n (%) | 8 (72) | 6 (50) | 6 (54) | 0.51 |
| Weight, Kg | 47.9 (5.8) | 50.3 (6.2) | 51 (7.1) | 0.47 |
| Temperature, °C | 38.0 (0.8) | 37.8 (0.7) | 37.4 (0.8) | 0.19 |
| Arterial pH | 7.51 (0.03) | 7.49 (0.02) | 7.49 (0.02) | 0.36 |
| Arterial Lactate, mmol/L | 1.3 (0.3) | 1.1 (0.2) | 1.5 (0.4) | 0.06 |
| Arterial Glucose, g/dl | 112.5 (31.7) | 97.0 (16.3) | 106.8 (13.1) | 0.24 |
| P/F | 435.7 (57.3) | 454.3 (68.0) | 482.4 (69.7) | 0.25 |
| PaCO<sub>2</sub>, mmHg | 41.6 (3.1) | 43.0 (1.7) | 41.2 (4.0) | 0.34 |
| Na<sup>+</sup>, mmol/L | 133.4 (2.37) | 134.6 (3.5) | 135.9 (5.3) | 0.36 |
| K<sup>+</sup>, mmol/L | 3.8 (0.3) | 3.7 (0.1) | 3.7 (0.3) | 0.27 |
| Cl<sup>-</sup>, mmol/L | 100.7 (1.5) | 101.8 (2.2) | 103.2 (5.0) | 0.21 |
| Base Excess, mEq/L | 9.0 (3.0) | 8.5 (1.1) | 7.7 (2.3) | 0.45 |
| Osmolarity, osm/L | 268.5 (3.2) | 269.7 (6.9) | 272.6 (10.2) | 0.43 |
| MAP, mmHg | 83 (12) | 89 (12) | 87 (11) | 0.57 |
| HR, bpm | 87 (14) | 105 (29) | 100 (22) | 0.22 |
| CVP, mmHg | 7.3 (3.2) | 6.6 (3.2) | 8.4 (3.3) | 0.47 |
| SvO<sub>2</sub>, % | 60.8 (7.9) | 63.3 (7.3) | 68.2 (6.9) | 0.15 |
| CO, L/min | 5.8 (1.1) | 6.0 (1.3) | 5.3 (0.5) | 0.28 |
| PPV, % | 11.4 (1.8) | 11.6 (3.5) | 12.8 (2.7) | 0.50 |
| PCWP, mmHg | 10.5 (2.9) | 9.5 (3.1) | 11.5 (3.1) | 0.29 |
| CPO, Watt | 0.8 (0.3) | 1.1 (0.3) | 0.9 (0.1) | 0.11 |
| DO<sub>2</sub>, ml/min | 624.5 (180.5) | 688.7 (193.3) | 608.7 (77.3) | 0.48 |
| VO<sub>2</sub>, ml/min | 245.7 (93.5) | 237.2 (65.7) | 190.2 (74.6) | 0.23 |
| OER, % | 39.4 (6.3) | 36.7 (8.6) | 31.0 (11.1) | 0.09 |
| Diuresis, ml | 600 (256-825) | 600 (300-700) | 600 (500-700) | 0.95 |
| ICP, mmHg | 8.0 (6.1-10.3) | 12.6 (6.8-14.4) | 10.1 (7.0-12.8) | 0.31 |
| PbtO<sub>2</sub>, mmHg | 33.2 (31.9 - 38.3) | 32.5 (25.7 - 41.6) | 40.1 (24.8 - 42) | 0.40 |
| Hemoglobin, g/dl | 8.9 (0.7) | 8.4 (0.7) | 8.4 (0.6) | 0.20 |

P/F = ratio between partial arterial oxygen pressure and fraction of inspired oxygen; PaCO<sub>2</sub> = arterial partial pressure of carbon dioxide; MAP= mean arterial pressure; HR, heart rate; CVP= central venous pressure; SvO<sub>2</sub>= mixed venous oxygen saturation; CO= cardiac output; PPV= pulse pressure variation; PCWP= pulmonary capillary wedge pressure; CPO= cardiac power output; DO<sub>2</sub>= oxygen delivery; VO<sub>2</sub>= oxygen consumption; OER= oxygen extraction ratio; ICP= intracranial pressure; PbtO<sub>2</sub>= brain tissue oxygen pressure;

### Cardiopulmonary Resuscitation

CPR lasted 300 (300-360) seconds in all groups. There was no significant difference between groups in maximal end-tidal CO_2_ (etCO_2_) over the first minute, the required number of countershocks, the epinephrine doses administered and the incidence of arrhythmia during the first 30 min after ROSC. A detailed report is presented in **Table S3**.

### Physiological and metabolic variables

After ROSC, arterial pH was higher in the treated groups than in the control group (p<0.001), as were the arterial lactate (p<0.001), sodium concentrations (p<0.001) and arterial plasmatic osmolarity (p<0.001), while potassium values were lower (p= 0.004; **Figure 3**). Arterial glucose levels did not differ between groups; only one animal required 10 units of intravenous insulin for hyperglycemia, two hours after ROSC.

**Figure 3:**
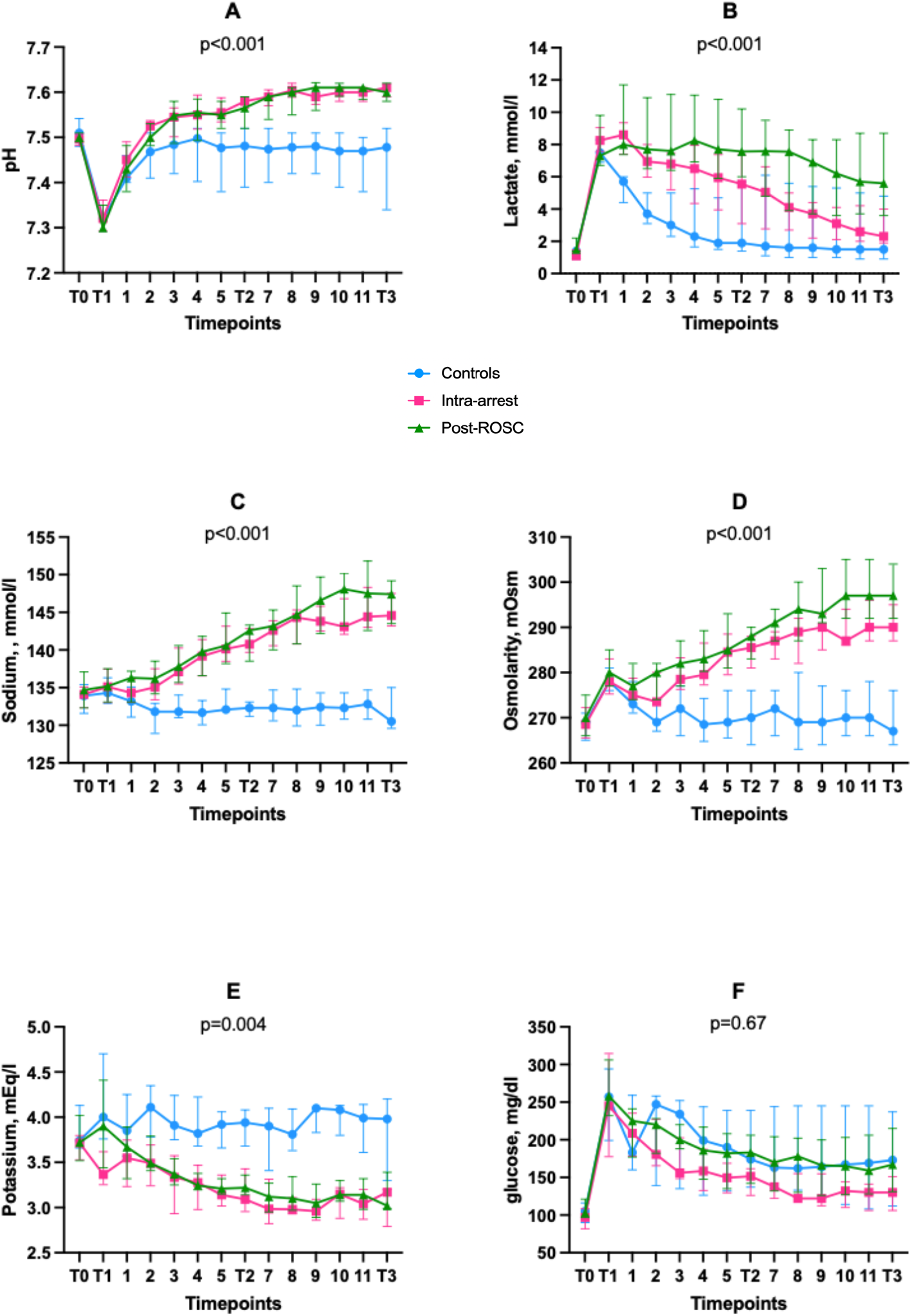
Time-course of main physiological variables and arterial electrolytes concentrations. A= Arterial pH; B= Arterial Lactate; C= Sodium; D= Osmolarity; E= Potassium; F= Arterial Glucose. Medians and bars interquartile ranges (25^th^-75^th^).

Arterio-jugular difference in glucose were lower and arterio-jugular difference in lactate levels higher in both treated groups compared to controls (p<0.001 and p<0.001, respectively - **Figure S1**). Despite a different trajectory, PaCO_2_ remained within the desired range and was similar in all groups (p= 0.02 for interaction, p=0.07 for groups).

### Hemodynamic variables

significant lower norepinephrine doses were required to maintain the target MAP ≥ 65 mmHg in the treated groups than in the controls (p= 0.005; **Figure 4**), while other hemodynamic variables, including cardiac output, central venous pressure, heart rate, pulse pressure variation and pulmonary artery wedge pressure were similar (**Table S4**). Despite a similar total fluid balance among groups (p=0.36), urine output was greater in treated groups than in controls (p<0.01, **Figure S2**).

**Figure 4:**
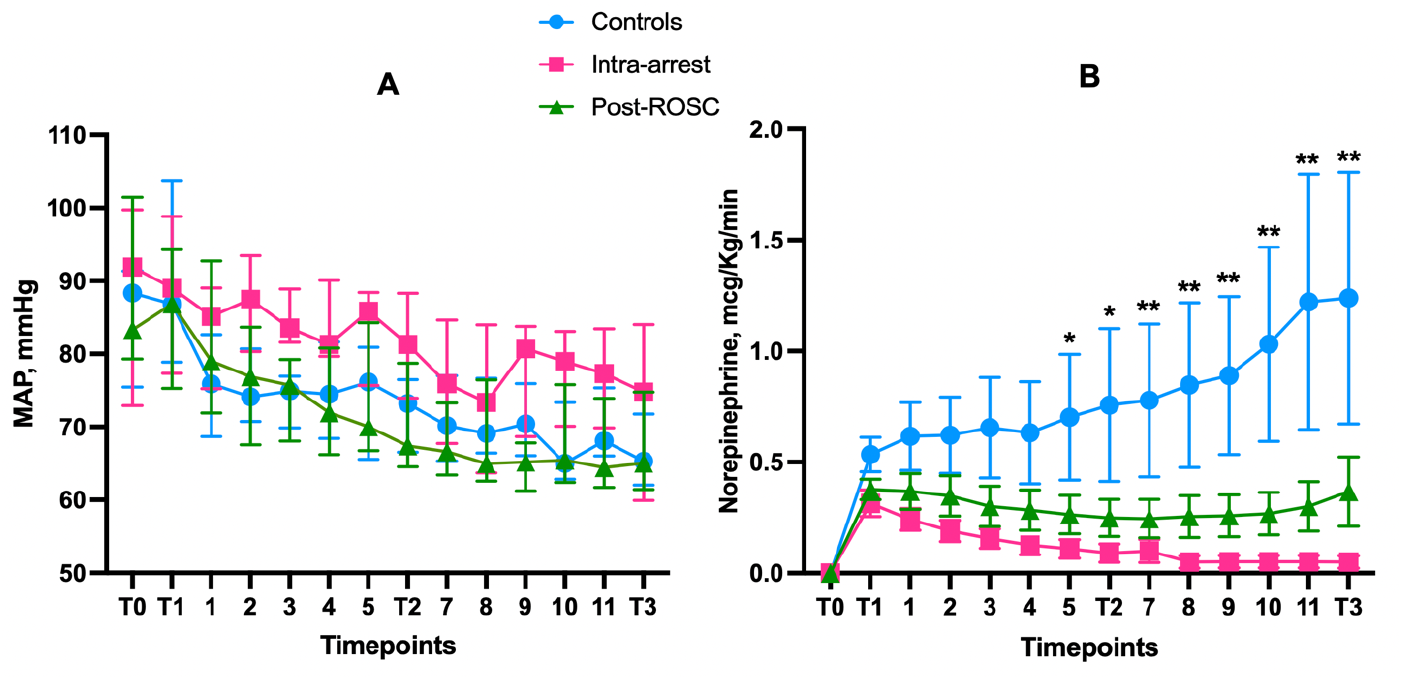
mean arterial pressure (median with interquartile range, A) and norepinephrine requirements (B) in the three groups (means and SEM; *: significantly different comparison Intra-arrest group and Controls; ** significantly different comparison between both HSL groups and Controls).

### Multimodal neuromonitoring

ICP, cerebral perfusion pressure and PbtO_2_, remained within the normal range for most of measurements with no difference among groups. No increase in the regional CBF was observed over time by laser Doppler between groups (**Figure S3**).

Cerebral lactate levels increased in the treated groups in the first hours after ROSC, when compared to Controls (p=0.004), while cerebral glucose, pyruvate, lactate to pyruvate ratio (LPR), glutamate and glycerol levels were similar among groups over time. (**Figure S4**).

### Plasma biomarkers

Circulating levels of troponin-I were lower in the treated groups than in the controls (**Figure 5A**), at both T2 and T3. No differences were identified among most of other measured variables, except for lactate dehydrogenases and alkaline phosphatases, which were lower in the treated groups in both T2 and T3 compared to Controls (**Figure S5**).

**Figure 5:**
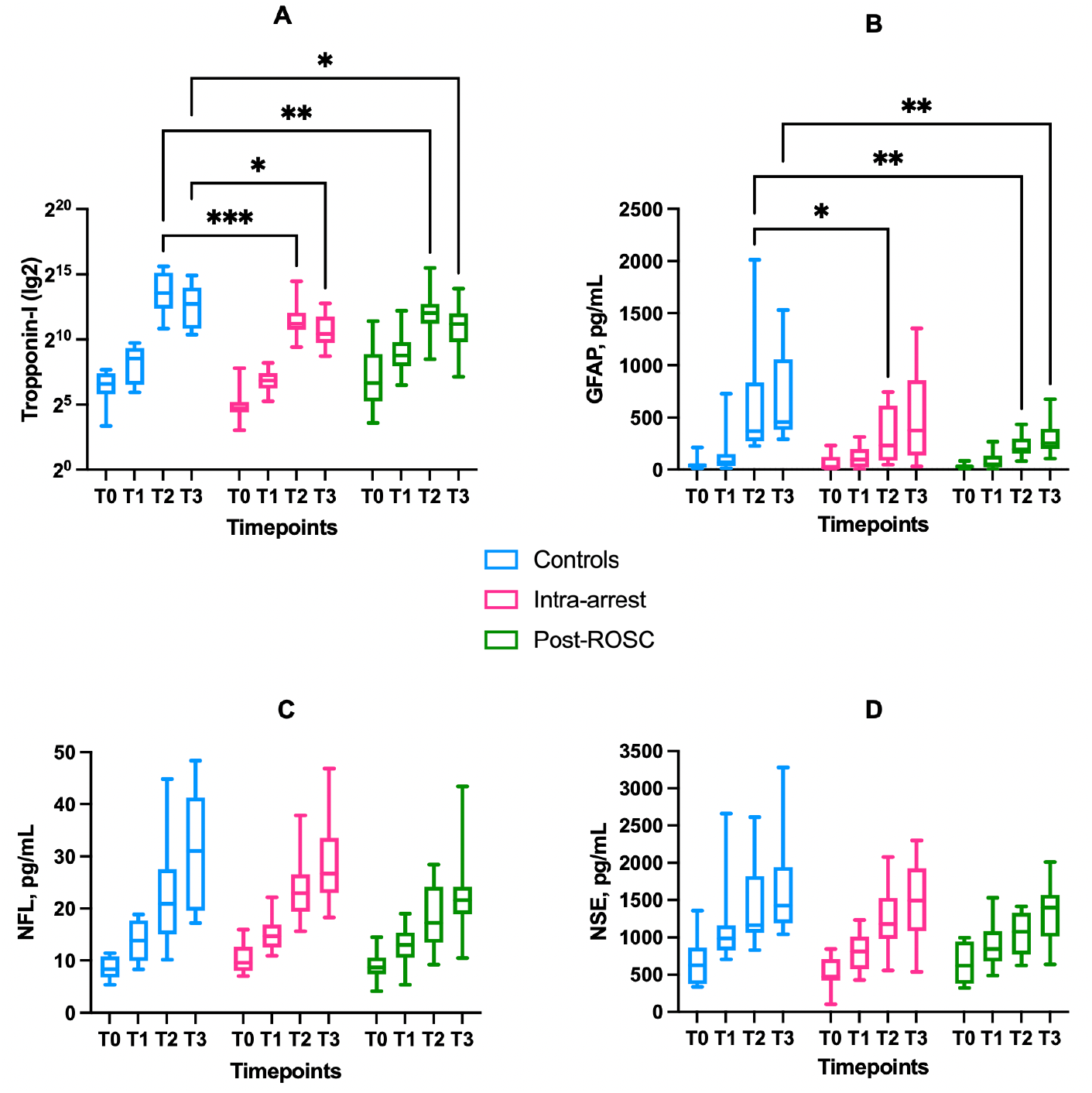
cerebral and cardiac plasmatic biomarkers. A= troponin I; B= glial fibrillary acid protein; C= neurofilament light chain; D= neuron-specific enolase. Boxes represent median values and interquartile ranges (25^th^-75^th^), whiskers extend between minimum and maximum values. * p<0.05; ** p<0.01; *** p<0.001.

Circulating GFAP levels were lower in the treated groups than in controls at T2 and T3, while NFL and NSE values tended to be also lower in the treated groups, although this difference did not achieve statistical significance (p=0.14 and p=0.39, **Figure 5C-D**).

### sEEG

There was a different trajectory over time of the mean amplitude and mean standard deviation between controls and pooled treated groups between T2 and T3 (p<0.001 for interaction in both cases), but not between T1 and T2 (p=0.86 and p=0.95, respectively), whereas a different trajectory was detected in mean kurtosis and mean skewness both between T1 and T2 and between T2 and T3 (p<0.001 for all interactions) (**Figure 6, S6**). At T3, there was a higher proportion of suppressed background or suppression-burst patterns in the Control groups when compared to the treated groups (n=8/22; 100 vs 63.6%, p=0.046) (**Figure S7**). No animal presented a status epilepticus at T3.

**Figure 6:**
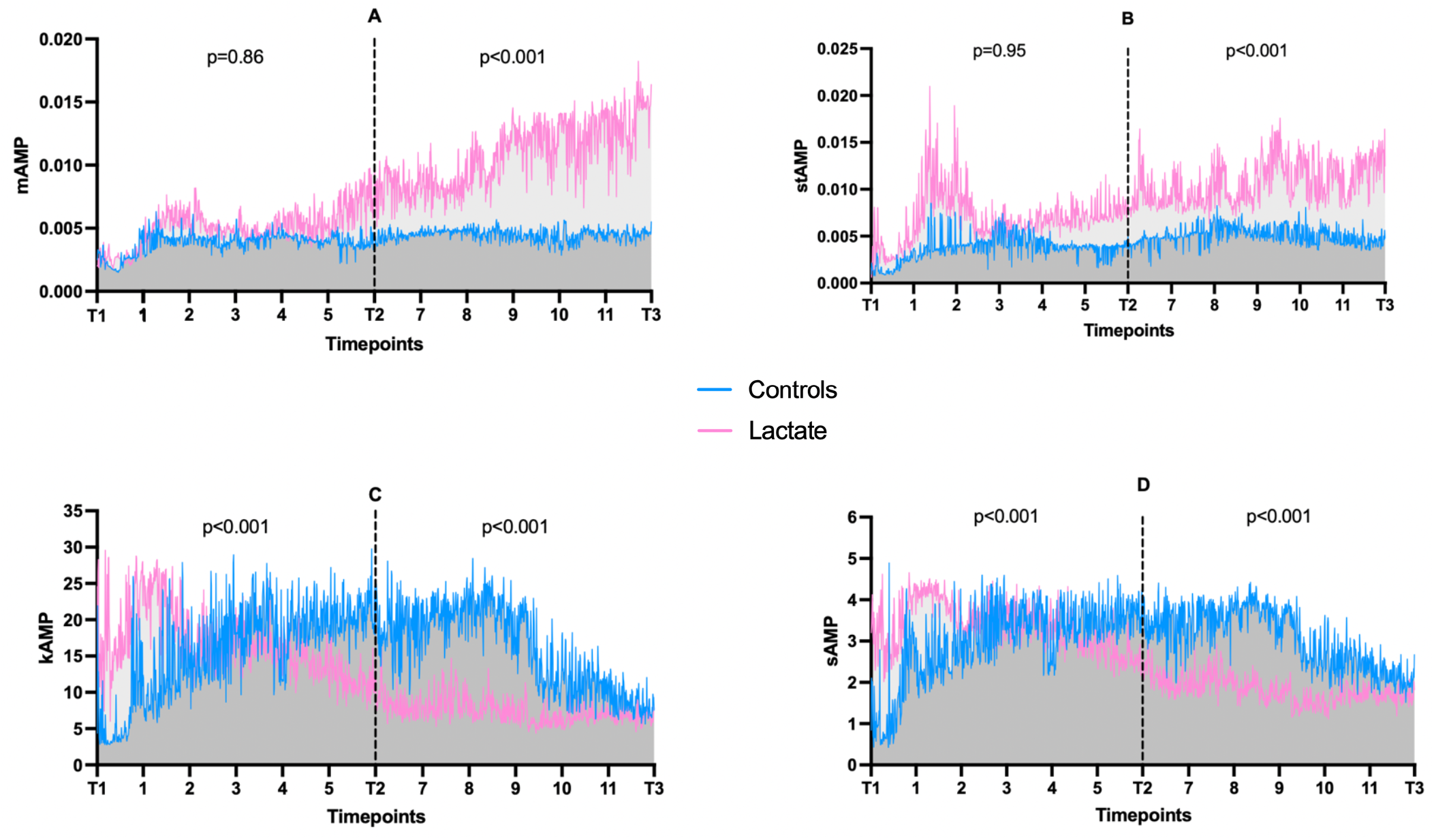
EEG evolution; A= mean amplitude; B= mean standard deviation; C= mean kurtosis; D= mean skewness. Lines connects median values over time, p values refer to interaction (time x group).

### Gene Expression

A reduced expression of HO-1 was found in both treated groups (p<0.01, **Figure 7A**) and PECAM-1 was lower in the Post-ROSC group compared to Controls (p=0.01) but not in the Intra-arrest group (p=0.05, **Figure 7B**). Apoptosis-related CASP-3 and CASP 8 was lower in the Post-ROSC group compared to Controls (p<0.01 and p=0.04, **Figure 7C**) No statistically significant differences between groups for the expression of MAP2, CD11ß and GFAP were found (**Figure 7D-G** and **Table S2**).

**Figure 7:**
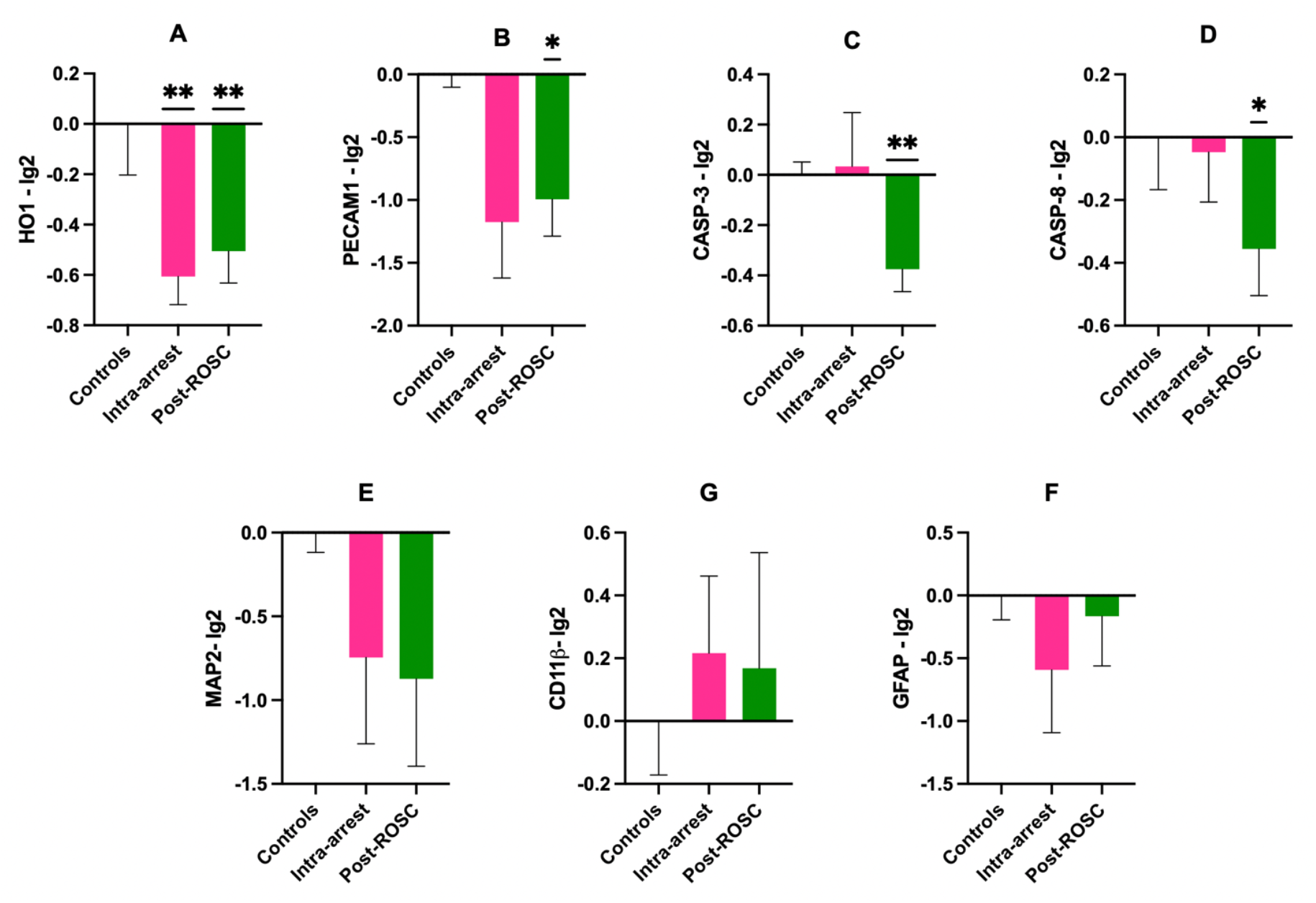
Gene expression in parietal cortex (Wilcoxon test). A= HO-1; B= PECAM1; C= CASP-3; D= CASP-8; E= MAP2; F= GFAP; G= CD11ß. Mean values SEM. For Controls n=4, Intra-arrest n= 5; Post-ROSC n=7. * p< 0.05; ** p < 0.01 versus controls.

## Discussion

Our findings demonstrate that HSL infusion during CPR and/or early after ROSC: A-reduces norepinephrine requirement in the first 12 hours after ROSC; B-mitigate brain and cardiac injury as indicated by a decrease in injury-related biomarkers and a reduced cortical expression of brain injury-related genes and C- was associated with a different cerebral activity, as pointed out by a different EEG evolution. Our experimental model was designed to achieve the maximum tolerated HSL infusion; despite the reduction of the infusion rate in the second half of the observation period due to safety limits, the treatment remained feasible for the entire duration of the experiment.

Along with a concurrently independent trial our study represents the first evidence of the application of HSL solution in the context of CA. Our findings are in line with the study by Stevin et al. [17], which showed in rabbits that the infusion of molar HSL for 120 minutes after ROSC increased the proportion of animals with pupillary reactivity at 2 h, reduced biomarkers of brain injury (astrocyte lysis marker S100β) improved hemodynamics (MAP, CO and left ventricular surface shortening fraction), and improved brain mitochondrial function. Similar to our results, HSL had a detectable effect on lactatemia, natremia and kaliemia between groups, and pH tended to increase in the treated group. Differently from our experiment, animals were treated with a higher HSL dose (i.e., 80 μmol/Kg/min vs 30 μmol/Kg/min) for a shorter period of time (i.e., 2 vs 12 hours) and the origin of CA (i.e., asphyxial vs electrical) and CPR techniques differed between our experimental designs. Furthermore, animals were not resuscitated aiming a minimal threshold in MAP and no target temperature management was used: by implementing these, we have limited the possible confounding effect of a different CPP and coronary perfusion pressure between groups when interpreting the organ damage. As such, the demonstration of short-term safety and the reproducibility of some benefits in two different preclinical models should pave the way to a clinical study assessing some physiological effects of HSL in this setting. In another trial, Miclescu et al compared HSL (0.63M), normal saline and hypertonic saline solution combined with methylene blue in a pig model of cardiac arrest.

Although no group received hypertonic sodium lactate as intervention, and the observation period lasted only 4 hours, in the methylene blue-sodium lactate group a lower concentration of CK-MB and Troponin I was observed compared to methylene blue-normal saline group as well as an alkalinizing effect in the HSL-methylene blue group. In contrast, no difference in plasmatic levels of protein S100β were found at 4 hours [18].

Treated animals needed less vasopressor support and had a greater urine output despite similar fluid requirements. Possibly, the rise of the pH contributed to ameliorate the effects of both endogenous and exogenous catecholamines [22]. On the other hand, despite a significant reduction of the myocardial injury as of a lower plasmatic level troponin-I at 6h and 12h in treated groups, no differences in cardiac output nor cardiac power output were observed. Previous studies suggested that lactate infusion could decrease myocardial injury after ischemia-reperfusion [23, 24]. One can reasonably hypothesize that the cardiac reserve of these previously healthy pigs was sufficient in condition of deep anesthesia and hypothermia to allow hemodynamic tolerance to the initial insult. It remains possible that, after discontinuation of TTM and sedation, with an increasing in systemic and myocardial VO_2_, myocardial dysfunction could become of clinical significance. Importantly, no significant differences were found between the Intra-arrest and the Post-ROSC groups, suggesting that a clinical trial could easily try to implement this therapy only in patients have a sustained and stable circulation.

After CA, we found a rapid increase of plasmatic GFAP levels, consistent with observations in patients [25]. Of note, a significant reduction in plasmatic GFAP was observed in HSL groups at 6 and 12 h post CA. While traditionally considered a marker of astrogliosis, blood GFAP levels have been suggested to reflect disruption of the blood brain barrier, both in acute CNS injury [26]and in highly inflammatory conditions such as COVID-19 [27] Indeed, GFAP is a key constituent of the astrocytic endfeets enveloping the blood brain barrier (BBB), which when damaged may release GFAP directly in the bloodstream. A reduction in BBB damage following treatment may thus underpin the neuroprotective effects of HSL. Although only NSE is recommended in clinical practice as outcome predictor after CA [28], other biomarkers have been suggested and appears to be more accurate [29]. To the best of our knowledge, this is the first large animal study in which NFL has been measured after CA after 12h observation period, although no significant differences were found between groups. Differently from what observed from GFAP, we did not see a statistically significant effect of NFL and NSE levels, although there is a trend towards decrease in the levels of the treated groups. While GFAP rapidly increases into the blood of patients with CA [29], NFL and NSE levels peak between 24 and 72 hours after ROSC [30,31]. As such, it is possible that with a prolonged observation period (i.e., 24-48 hours), different trajectories of these biomarkers (i.e., NSE and NFL) might have been observed among study groups.

Although no significant differences in ICP, PbtO_2_ and regional CBF were observed when HSL was infused, ICP and PbtO_2_ remained within the normal range, thus limiting the potential impact of HSL on these variables. Nonetheless, heterogeneous ICP values have been reported in CA survivors and we cannot infer on the possible utility of hypertonic infusions in these subjects [32–34]. Moreover, the increase in pH and decreased cerebral temperature in treated animals might have, theoretically, increased the hemoglobin affinity for oxygen by shifting the Hb dissociation curve to the left, reducing the interstitial oxygen availability and, therefore, absolute PbtO_2_ values. Despite its metabolic effect, no difference in cardiac output or cerebral perfusion pressure were detected, suggesting that the overall cerebral DO_2_ didn’t differ during the observation period. Despite cerebral metabolites explored with CMD were similar in the groups, except for an increase in cerebral lactate, the arterio-jugular difference of lactate and glucose suggested that an increased uptake of lactate with a glucose sparing effect was present at some timepoints in the treated groups, thus underlying the possible metabolic shift between different substrates in case of acute injury. One could also argue that the use of deep sedation and TTM decreased cerebral metabolic rate [35,36] and possibly underestimated the impact of HSL on cerebral function. Although the increased EEG activity could not directly relate to a better prognosis, intracerebral EEG recordings suggested an earlier recovery of the background rhythm, which might be secondary to a more rapid restoration of cerebral metabolism or perfusion. HSL has been proved capable to improve mitochondrial function [17], and we have found a reduced gene expression related to oxidative stress (HO1), apoptosis (CASP-3 and CASP-8) and endothelial function. There were no significant differences for neuronal, microglial, and astrocytic related genes. While plasmatic GFAP, NFL and NSE levels are indicative of the damage to existing cerebral cells at the time of injury, gene expression analyses reflect cell reaction after injury. The lack of treatment effects on these parameters suggests that later time points should be investigated to observe possible reparative/compensative effects induced by HSL. Nevertheless, all together, our data suggests that the favorable effects on brain damage of HSL infusion in CA are related to multiple mechanisms rather than a single one.

Our study has several strengths: our model of CA allowed at the same time a good survival rate and a severe post-resuscitation vasoplegia and cerebral damage. Moreover, many different physiological parameters, including multimodal neuromonitoring, intracranial EEG and continuous cardiac output monitoring were continuously collected. Third, our observations were prolonged over a relatively long period after-ROSC, intercepting physiological modifications produced by both primary (i.e., minutes) and secondary (i.e., hours) injuries after hypoxic ischemic brain injury.

Our study also has several limitations: first, we selected an arbitrary dosage of the intra-arrest bolus of HSL, and this might have been insufficient to elicit a sensible biological effect. Nevertheless, it is possible that a different experimental design with longer CA duration would increase the possibility for a clinical impact of the intra-arrest bolus of HSL. Second, swine present differences in lactate metabolism compared to humans and this might limit the applicability of the present results to other species. Nevertheless, the swine cardiovascular system and brain architecture closely resemble humans, providing a very reliable experimental model. Third, no ultrastructural and morphological analyses were performed, as well as no cardiac tissue analysis. Fourth, despite different areas of the brain might exhibit different susceptibility to hypoxic ischemic injury, the gene expression was explored only in the parietal cortex. Fifth, regional multimodal neuromonitoring was used, and therefore caution should be used in generalizing our results to the brain parenchyma as a whole.

However, this approach is derived from current neurocritical care practice in humans; additionally, as the hypoxic event is rather homogenous, regional monitoring should still be able to detect relevant change in main physiological variables. Sixth, no additional hemodynamic monitoring (i.e., pressure/volume curves or echocardiography) was performed. Nonetheless, we aimed to minimize the possibility to induce ventricular arrhythmia in the post-resuscitation phase, and the ultrasound window was limited by the ventral position of the animal after ROSC.

## Conclusions

In this experimental large animal model of CA, HSL, either started during CPR or after ROSC, was associated with reduced norepinephrine requirements and lower levels of circulating biomarkers of cardiac and brain injury.

## Acknowledgements

FA, FST, JC, NG and JLV conceived the protocol; FA, FS and LP conducted the experiment procedures; FA, IL, EC, FP and ERZ performed cerebral biomarker tests and genic expression analyses; FA, HN and EGB elaborated the statistical analysis of the data. NG and LF analyzed the EEG trace; FA wrote a first draft of the manuscript. All authors provided substantial intellectual contribution, participated in the modification of the first draft and approved the final version of this manuscript.

## Source of founding

Dr Filippo Annoni was supported by a research grant during the period of this study by Fonds Erasme pour la Recherche Médicale (2018-2020, and an additional semester during the year 2020-2021). Dr Filippo Annoni has received support for research mobility in 2021 by the Fonds de la Recherche Scientifique (FRS-FNRS). Pr Fabio Silvio Taccone was also supported by a research grant during the period of this study by Fonds Erasme.

## References

1. Perkins DG, Grasner JT, Semeraro F, Olasveengen T, Soar J et al. European Resuscitation Council Guidelines 2021: executive summary. Resuscitation 2021; 161: 1–60.

2. Grasner JT, Herlitz J, Tjelmeland I BM, Wnent J, Masterson S et al. European Resuscitation Council Guidelines 2021: epidemiology of cardiac arrest in Europe. Resuscitation 2021; 161: 1–60.

3. Lemiale V, Dumas F, Mongardon N, Giovanetti O, Charpentier J, Chiche JD. Intensive care unit mortality after cardiac arrest: the relative contribution of shock and brain injury in a large cohort. Intensive Care Med 2013; 39:1972-1980.

4. Taccone FS, Crippa IA, Dell’Anna AM, Scolletta S. Neuroprotective strategies and neuroprognostication after cardiac arrest. Best Pract Res Clin Anaesthesiol 2015; 29: 451–464

5. Lascarrou JB, Merdji H, Le Gouge A et al. Target temperature management for cardiac arrest with no shockable rhythm. N Engl J Med 2019; 381(24): 2327–2337.

6. Taccone FS, Hollenberg J, Forsberg S, Truhlar A, Jonsson M et al. Effect of intra-arrest trans-nasal evaporative cooling in out-of-hospital cardiac arrest: a pooled individual partecipand data analysis. Crit Care 2021; 25: 198.

7. Dankiewicz J, Cronberg T, Lilja G, Jackobsen J, Levin H et al. Hypothermia versus normothermia after out-of-hospital cardiac arrest. NEJM 2021; 384: 2283–2294.

8. Ichai C, Armando G, Orban JC, Berthier F, Rami L et al. Sodium lactate versus mannitol in the treatment of intracranial hypertensive episodes in severe traumatic brain-injured patients. Intensive Care Med. 2009; 35:471–479.

9. Ichai C, Payen JF, Orban JC, Quintard H, Roth H, Legrand R, Francony J, Leverve XM. Half-molar sodium lactate infusion to prevent intracranial hypertensive episodes in severe traumatic brain injured patients: A randomized controlled trial. Intensive Care Med. 2013; 39: 1413–1422.

10. Quintard H, Patet C, Zerlauth JB, Suys T, Bouzat P, Pellerin L, Meuli R, Magistretti PJ, Oddo M. Improvement of neuroenergetics by hypertonic lactate therapy in patients with traumatic brain injury is dependent on baseline cerebral lactate/pyruvate ratio. J. Neurotrauma 2016; 33: 681–687.

11. Carteron L, Solari D, Patet C, Quintard H, Miroz JP, Bloch J, Daniel RT, Hirt L, Eckert P, Magistretti PJ et al. Hypertonic lactate to improve cerebral perfusion and glucose availability after acute brain injury. Crit. Care Med. 2018; 46: 1649–1655.

12. Bouzat P, Sala M, Suys T. Cerebral metabolic effect of exogenous lactate supplementation on the injured human brain. Intensive Care Med. 2014; 40: 412–421.

13. Bernini A, Miroz JP, Abed-Maillard S, Favre E, Iaquaniello C et al. Hypertonic lactate for the treatment of intracranial hypertension in patients with acute brain injury. Sci Rep 2022; 12: 3035.

14. Annoni F, Peluso L, Gouvea Bogossian E, Creteur J, Zanier ER and Taccone FS. Brain protection after anoxic brain injury: is lactate supplementation useful? Cells 2021; 10: 1714.

15. Nalos M, Leverve XM, Huang SJ, Weisbrodt L, Parkin R, et al. A Half-molar sodium lactate infusion improves cardiac performance in acute hearth failure: A pilot randomised controlled clinical trial. Crit Care 2014; 18: 1–9.

16. Nalos M, Kholodniak E, Smith L, Orde S, Ting I, Slama M, Seppelt J, McLean AS, Huang S. The comparative effects of 3% saline and 0.5M sodium lactate on cardiac function: A randomised, crossover study in volunteers. Crit Care Resusc 2018; 20: 124–130.

17. Stevic N, Argaud L, Loufouat J, Kreitmann L, Desmurs L, Ovize M, Bidaux G, cour M. Molar sodium lactate attenuates the severity of postcardiac arrest syndrome: a preclinical study. Crit Care Med 2022; 50(1): e71–e79.

18. Micelscu A, Basu S, Wiklund L. Cardio-cerebral and metabolic effects of methylene blue in hypertonic sodium lactate during experimental cardiopulmonary resuscitation. Resuscitation 2007; 75(1): 88–97.

19. Annoni F, Peluso L, Hirai LA, Babini G, Khaldi A, Herpain A, Pitisci L, Ferlini L, Garcia B, Taccone FS, Creteur J, Su F. A comprehensive neuromonitoring approach in a large animal model of cardiac arrest. Anim Models Exp Med 2022; 00: 1–5.

20. Westhall E, Rossetti AO, van Rootselaar AF, Wesenberg Kjaer T, Horn J et al. Standardized EEG interpretation accurately predicts prognosis after cardiac arrest. Neurology 2016; 86(16): 1482–1490.

21. Babini G, Grassi L, Russo I, Novelli D, Boccardo A et al. Duration of untreated cardiac arrest and clinical relevance of animal experiments: the relationship between the “no-flow” duration and the severity of post-cardiac arrest syndrome in a porcine model. SHOCK 2018; 49: 205–212.

22. Weil MH, Houle DB, Brown EB and Campbell GS. Vasopressor agents. Influence of acidosis on cardiac and vascular resistance. California Medicine 1958; 88(6): 437–440.

23. Koyama T, Shibata M, Moritani K. Ischemic postconditioning with lactate-enriched blood in patients with acute myocardial infarction. Cardiology 2013; 125: 92–93.

24. Koyama T. Lactate Ringer’s solution for preventing myocardial reperfusion injury. IJC Heart & vasculature 2017; 15:1–8.

25. Whiersaari L, Reinikainen M, Furlan R, Mandelli A, Vaahersalo J et al. Neurofilament light chain compared to neuron-specific enolase as a predictor of unfavourable outcome after out-of-hospital cardiac arrest. Resuscitation 2022; 174: 1–8.

26. Graham NSN, Zimmerman K A, Moro F, Helsgrave A, Maillard SA, Bernine A et al. Axonal marker neurofilament light predicts long-term outcomes and progressive neurodegeneration after traumatic brain injury. Sci Trans Med 2021; 13(623): eabg9922.

27. Bonetto V, Pasetto L, Lisi I, Cabonara M, Zangari R, Ferrari E et al. Markers of blood-brain barrier disruption increase early ane persistently in COVID-19 patients with neurological manifestations. Front Immunol 2022; 13: 1070379.

28. Nolan J, Sandroni C, Bottiger BW, Cariou A, Cronberg T et al. European Resuscitation Council and European Society of Intensive Care Medicine Guidelines 2021: post-resuscitation care. Resuscitation 2021; 161: 220–269.

29. Kaneko T, Kasaoka S, Miyauchi T, Fujita M, Oda Y et al. Serum glial fibrillary acid protein as predictive biomarker of neurological outcome after cardiac arrest. Resuscitation 2009; 80: 790–794.

30. Harel A, Marten K, Salla N, Lasse V. Biomarkers of traumatic brain injury, temporal changes in body fluuids. eNeuro 2016; 3(6): e0294

31. Whihersaari L, Ashton NJ, Reinikainen M, Pettila V, Hastbacka J, et al. Neurofilament light as an outcome predictor after cardiac arrest: a post-hoc analysis of the COMACARE trial. Intensive Care Med 2021; 47: 39–48.

32. Sekhon MS, Griesdale DE. Individualized perfusion targets in hypoxic ischemic brain injury after cardiac arrest. Crit Care 2017; 21: 259–263.

33. Sekhon MS, Gooderham P, Manon DK, Brasher PMA, Foster D et al. The burden of brain hypoxia and optimal mean arterial pressure in patients with hypoxic ischemic brain injury after cardiac arrest. Crit Care Med 2019; 47 (7): 960–969.

34. Sekhon MS, Griesdale DE, Ainslie PN, Gooderham P, Foster D et al. Intracranial pressure and compliance in hypoxic ischemic brain injury patients after cardiac arrest. Resuscitation 2019; 141: 96–103.

35. Yager JY and Asselin J. Effect of mild hypothermia on cerebral energy metabolism during the evolution of hypoxic-ischemic brain damage in the immature rat. Stroke 1996; 27: 919–926.

36. Slupe AM and Kirsch JR. Effect of anesthesia on cerebral blood flow, metabolism, and neuroprotection. J Cereb Blod Flow Metab 2018;38(12): 2192–2208.

